# Using variable data independent acquisition for capillary electrophoresis-based untargeted metabolomics

**DOI:** 10.1101/2024.01.17.576157

**Authors:** Saki Kiuchi, Yasuhiro Otoguro, Tomoaki Nitta, Mi Hwa Chung, Taiki Nakaya, Yuki Matsuzawa, Katsuya Oobuchi, Kazunori Sasaki, Hiroyuki Yamamoto, Hiroshi Tsugawa

**Affiliations:** Department of Biotechnology and Life Science, Tokyo University of Agriculture and Technology, 2-24-16 Naka-cho, Koganei, Tokyo 184-8588, Japan; Human Metabolome Technologies Inc., 246-2 Mizukami, Kakuganji, Tsuruoka, Yamagata 997-0052, Japan; Tsumura Kampo Laboratories, Tsumura&Co, Ami, Ibaraki 300-1192, Japan; RIKEN Center for Integrative Medical Sciences, 1-7-22 Suehiro-cho, Tsurumi-ku, Yokohama, Kanagawa 230-0045, Japan; RIKEN Center for Sustainable Resource Science, 1-7-22 Suehiro-cho, Tsurumi-ku, Yokohama, Kanagawa 230-0045, Japan; Molecular and Cellular Epigenetics Laboratory, Graduate School of Medical Life Science, Yokohama City University, 1-7-29 Suehiro-cho, Tsurumi-ku, Yokohama, Kanagawa 230-0045, Japan

## Abstract

Capillary electrophoresis coupled with tandem mass spectrometry (CE-MS/MS) offers advantages in peak capacity and sensitivity for metabolic profiling, owing to the electroosmotic flow-based separation. However, the utilization of data-independent MS/MS acquisition (DIA) is restricted due to the absence of an optimal procedure for analytical chemistry and its related informatics framework. We assessed the mass spectral quality using two DIA techniques, namely, all-ion fragmentation (AIF) and variable DIA (vDIA), to isolate 60∼800 Da precursor ions with respect to annotation rates. Our findings indicate that vDIA, coupled with the updated MS-DIAL chromatogram deconvolution algorithm, yields higher spectral matching scores and annotation rates compared to AIF. Additionally, we evaluated a linear migration time (MT) correction method using internal standards to accurately align chromatographic peaks in a dataset. After the correction, the peaks exhibited less than 0.1 min MT drifts, a difference mostly equivalent to that of conventional reverse-phase liquid chromatography techniques. Moreover, we conducted MT prediction for metabolites recorded in mass spectral libraries and metabolite structure databases containing a total of 469,870 compounds, achieving an accuracy of less than 1.5 min root mean squares. Thus, our platform provides a peak annotation platform utilizing MT information, accurate precursor *m/z*, and the MS/MS spectrum recommended by the metabolomics standards initiative. Applying this procedure, we investigated metabolic alterations in lipopolysaccharide (LPS)-induced macrophages, characterizing 170 metabolites. Furthermore, we assigned metabolite information to unannotated peaks using an *in-silico* structure elucidation tool, MS-FINDER. The results were integrated into the nodes in the molecular spectrum network based on the MS/MS similarity score. Consequently, we identified a significantly increased amount of metabolites in the LPS-administration group, glycinamide ribonucleotide, not present in any spectral libraries. Additionally, we retrieved metabolites of false-negative hits in the initial spectral annotation procedure. Overall, our study underscores the potential of CE-MS/MS with DIA and computational mass spectrometry techniques for metabolic profiling.

---

One of the goals of metabolomics is to comprehensively characterize the metabolome in cellular systems. Capillary electrophoresis coupled with mass spectrometry (CE/MS) has been employed for the highly sensitive profiling of polar and charged compounds^1,2,3^. The CE-MS data obtained in this study reveal that CE/MS exhibits a narrower peak width (0.46 min on average) and a higher signal-to-noise ratio (*S/N*, approximately 2500 on average) compared to reversed-phase liquid chromatography MS (LC/MS) used in our previous study for untargeted lipidomics^4^, with an average peak width of 0.58 min and *S/N* of approximately 1000 (**Figures S1a, S1b**). This tendency can be attributed to the electroosmotic flow (EOF) in CE, which provides a nearly uniform velocity distribution and a lower buffer flow rate compared to the conditions used in LC^5^. Additionally, while the principle of CE focuses on polar/charged chemicals, 69% of the metabolites recorded in the human metabolome database (HMDB)^6^ were identified as ionizable molecules under the pH conditions used in CE-MS analysis (**Table S1, Figure S1c**). These findings indicate that CE/MS is a powerful approach for untargeted metabolomics.

The best practice for metabolite annotation in untargeted metabolomics involves similarity matching of migration time (MT), accurate precursor *m/z*, isotope ratio, and MS/MS spectrum with reference spectral libraries^7,8^. Obtaining comprehensive MS/MS and high reproducibility for MT can enhance confidence in peak annotation and expand metabolome coverage for global profiling. Data-independent MS/MS acquisition (DIA) has emerged as a powerful approach for untargeted metabolomics due to its capability to acquire all MS2 spectra and its high quantitative accuracy. With DIA-MS, all detected precursor ion peaks have MS/MS spectrum information, and the spectrum mixture is purified using the chromatogram deconvolution algorithm ^9,10,11^. However, the use of DIA coupled with chromatogram deconvolution has not been validated for untargeted metabolomics using CE-MS/MS. Additionally, large MT drift between samples often results in MT differences of more than 1.0 min, making peak annotation and alignment challenging^12,13,14^. In fact, high reproducibility of chromatographic peak elution time (less than 0.2 min between sample injections) increases confidence in peak alignments and reduces false peak calls in the alignment result. Thus, chromatographic peak adjustments using MT “signposts” with internal standards are recommended before the peak alignment function is performed.

In this study, we developed a platform for CE-MS/MS-based untargeted metabolomics, evaluating and optimizing three techniques—DIA, MT correction, and MT prediction—to enhance the confidence in data integration and peak annotation. Two DIA methods, all-ion fragmentation (AIF) and variable DIA (vDIA), were assessed for spectral quality and annotation rate. To handle vDIA data, we updated the MS-DIAL^4^ algorithm. We also examined the retention correction function of MS-DIAL to efficiently align chromatographic peaks among samples. Migration times of metabolite ion peaks were linearly corrected along the MT axis using internal standard compounds. Finally, we developed an MT prediction model using chemical descriptors from the PaDEL^15^ and ChemAxon (https://www.chemaxon.com) programs. Predicted MT information was incorporated into the mass spectral library with 44,257 molecules and the metabolite structure database with 446,485 molecules. For the MT prediction model, various machine learning methods were investigated using the Retip R package^16^. The platform was applied to untargeted hydrophilic metabolome profiling of lipopolysaccharide (LPS)-treated macrophage cells to comprehend metabolic alterations in an acute inflammatory process induced by bacteria. Our platform distinctly elucidates the metabolic rewiring of macrophages. Additionally, we demonstrated an integrated approach for unknown MS/MS spectral annotation. Nodes in molecular spectrum networking were further annotated using an *in-silico* structure elucidation tool, MS-FINDER^17^, with predicted MT values.

## MATERIALS AND METHODS

### Cell culture and treatment

RAW 264.7 cells were grown at 37 °C in DMEM (Dulbecco’s modified Eagle medium) supplemented with 10% EquaFETAL (Atlas Biologicals, Inc.) in a humidified 5% CO_2_ atmosphere. For sample preparation, the cells (5×10^5^ cells per well) were incubated in six-well plates (Thermo Fisher Scientific, Nunclon Delta) for 2 h before stimulation with 100 ng/mL lipopolysaccharide (from *E. coli* O55:B5, Sigma Aldrich), with four biological replicates per condition. At 1 h and 24 h after LPS stimulation, cell collection and metabolite extraction were performed according to a previously described protocol^18^. Briefly, the medium was removed and washed twice with 5% mannitol. After adding 800 μL of methanol, the solvent was incubated for 30 s, followed by the addition of 550 mL of Human Metabolome Technologies (HMT) internal standard solution 1 (Human Metabolome Technologies, Inc., Tsuruoka, Japan), with an additional 30-s incubation period. An aliquot of 1000 mL was transferred to a 1.5-mL tube and centrifuged at 4 °C and 2,300 g for 5 min. The supernatant (350 mL) was subjected to ultrafiltration to remove macromolecules. Thereafter, the filtrate was dried *in vacuo* and reconstituted in 50 mL of Milli-Q water for CE/MS analysis.

### Capillary Electrophoresis setting

Capillary electrophoresis was performed using a CE System 7100 (Agilent Technologies Inc., Santa Clara, CA, USA), and an Agilent 1260 isocratic HPLC pump was used to create the sheath flow. The sheath-flow rate was set at 10 μL/min. Metabolites were separated using a fused silica capillary (50 μm i.d. × 80 cm total length; Polymicro Technologies, Inc., Phoenix, AZ) with 50 mM ammonium acetate (pH 8.5) for anion analysis and 1 M formic acid for cation analysis (pH 1.8). The applied voltage for CE was set at 30 kV. The systems were controlled by MassHunter software version B.08.00 (Agilent Technologies Inc.).

### Mass Spectrometer setting

An Orbitrap mass spectrometer, Q Exactive Plus (Thermo Fisher Scientific Inc., Waltham, MA), was equipped with an ESI adapter^19^ developed in-house under the following conditions: sheath-gas flow, 5 L/min; dry-gas flow, 5.5 L/min; dry-gas temperature, 350 °C; interface voltage for anion analysis, 3.5 and 3.0 kV for primary and secondary electrodes, respectively; and interface voltage for cation analysis, −4.0 and −3.5 kV for primary and secondary electrodes, respectively. The mass spectrometer inlet capillary temperature was set at 100 °C for anion analysis and 50 °C for cation analysis. The magnitude of the S-lens RF level was set at 50 for cation analysis and 90 for anion analysis. The full scan ranges for cation and anion analyses were set to *m/z* 60–500 and *m/z* 70–800, respectively. The AIF parameters were set as follows: collision energy, 20 V; mass resolution, 70,000 at *m/z* 200. The parameters for the vDIA were set as follows: collision energy, 20 V; mass resolution, 17,500 at *m/z* 200. In the vDIA analysis, the precursor isolation window was set as described in **Table S2**, optimized by the ion signal density along with the migration time (**Figure S2**).

### Migration time prediction

A total of 13,677 descriptors were generated by the PaDEL-descriptor program^15^ and ChemAxon’s ‘cxcalc’ function (http://www.chemaxon.com). The ChemAxon cxcalc was utilized to generate six physicochemical properties, including formalcharge, chargedistribution, apka1, bpkja1, molecularpolarizability, and exactmass. Additionally, the ratio of exact mass to molecular polarizability property values was used as a chemical descriptor to describe *m/z*. Values with zero standard deviations were excluded. A total of 1,363 and 1,535 descriptors were used in the models for anion- and cation analyses, respectively. We excluded structures from our search space whose descriptors were not generated successfully by PaDeL or ChemAxon programs for unidentified reasons, resulting in the generation of chemical descriptors for 21,868 and 15,337 structures out of 35,635 and 23,409 compound records in the spectral database for cation and anion analysis, respectively, and for 349,218 structures out of 446,356 records in the MS-FINDER structure database. Machine learning was performed using the R software environment (version 4.2). The best machine learning model was selected using the Retip^16^ function with the generated descriptors. A total of 326 and 285 compounds were used as the training dataset for cation and anion analyses, respectively. The migration times were confirmed using the in-house standard compound database. The XGBoost model was selected as the best learner. The parameters were further optimized using the fit.xgboost() function of Retip, and the model accuracy was evaluated using statistical scores obtained from the get.score() function. Finally, the migration times for the structures recorded in the MS/MS library and the MS-FINDER structure database were predicted using the RT.spell() function.

### Data processing

Peak picking, chromatogram deconvolution, annotation, migration time correction, and peak alignment of the CE-MS/MS dataset were performed using MS-DIAL. Because the vDIA method uses a tricky scanning sequence, as described in **Figure S2**, the MS-DIAL program was updated to correctly recognize the MS1 and MS/MS frames. The migration time correction, i.e., data point correction, was performed using the migration time information of the internal standard compounds, including methionine sulfone (cation analysis) and camphor sulfonic acid. As the MS-DIAL program can import multiple spectral libraries with defined priority levels, three MSP spectral libraries were used for metabolite annotation. The most prioritized spectral library contains the migration time of metabolites confirmed by authentic standards, named “msp1.” The secondly prioritized spectral library contains the predicted migration time of metabolites generated by our machine learning model, named “msp2.” The last library, named “msp3,” has no MT information. The retention time tolerance for msp1 in annotation was set to 0.2 min, and the tolerances for msp2 were set to 1.48 min and 1.96 min for cation- and anion libraries, respectively. The tolerances were determined based on 95% confidence intervals in the machine-learning models. The following MS-DIAL parameters were changed from the default settings: minimum amplitude in peak detection, 10,000; retention time tolerance in peak alignment, 0.5 min. All the parameters used are described in **Table S3**.

### Statistics

Peak heights were corrected by the cell number and used for the statistical analysis below. The statistical significance for two-group comparison was calculated by the Mann-Whitney U test for each metabolite using the “exactRankTests” package of the R language (version 4.2.2), and *p*-values were adjusted using the Benjamini Hochberg method. Fold changes were also calculated, and the fold changes and *p*-values were used to create a volcano plot. Principal component analysis was performed using the “prcomp” function in the R environment, and auto scaling (also known as unit variance) was used for the data scaling method. Enrichment analysis was performed using MetaboAnalyst 5.0^20^ for the significantly changed metabolites between the two groups using the KEGG (Kyoto Encyclopedia of Genes and Genomes)^21^ metabolites set library.

### Metabolite annotation using MS-FINDER

The spectral data, including peak ID, precursor *m/z*, adduct type, migration time, and MS/MS in the alignment result, were exported in mat format from the MS-DIAL program. The query files were imported into MS-FINDER^17^. Metabolite annotation was performed with the following parameters: search option, formula prediction, and structure elucidation by *in silico* fragmenter; precursor ion option, precursor-oriented spectral search; mass tolerances for MS1 and MS2, 0.005 Da and 0.01 Da, respectively; data source, HMDB^5^. Additionally, the predicted MT values were imported for structure elucidation, and the retention time tolerances for cation and anion analyses were set to 1.48 min and 1.96 min, respectively. The tolerances were based on 95% confidence intervals (CIs).

### Construction of molecular spectrum network

The nodes and features of the MS/MS spectra were connected by the similarity of the product ion spectra based on the Bonanza algorithm^22^. When the two spectra were compared, the product ion peak was defined as a match if the difference in the product ion or neutral loss was within 0.05 Da. The base peak was normalized to 100, and product ions with both a relative abundance and an absolute abundance exceeding 0.1 and 50.0, respectively, were utilized to calculate scores after square-root scaling. The minimum peak match, cutoff score, and maximum edge number per node were set to 1, 0.8, and 5, respectively. The node and edge tables were generated using the MS-DIAL program. After the nodes and edge tables were created using the features of the macrophage cells’ experimental spectra, each of the experimental features was further connected with the features of the spectral libraries. The exported tables were imported into Cytoscape 3.9.1 program^23^. Authentic standard compounds were measured to validate the spectral annotation using MS-FINDER and molecular spectrum networking. Authentic standard compounds of 3-aminoadipic acid, 2-thio PAF (1-*O*-hexadecyl-2-deoxy-2-thio-*S*-acetyl-*sn*-glyceryl-3-phosphorylcholine), and *N*-acetylneuraminic acid were purchased from Fujifilm Wako (Tokyo, Japan), Santa Cruz Animal Health (Texas, US), and Tokyo Chemical Industry Co. (Tokyo, Japan). The standard compound was adjusted to a final concentration of 10 mM, and the solution, including the internal standards, was analyzed as described above.

## RESULTS AND DISCUSSION

### Workflow of CE-MS/MS-based untargeted analysis for comprehensive metabolome annotation

Here, we propose a procedure for untargeted metabolomics using CE-MS/MS (**Figure 1**). We prepared LPS-induced macrophage, RAW 264.7 cells for the evaluation of DIA, migration time correction, and metabolite annotation. The metabolite extracts of the cells after 1 h and 24 h of LPS administration were subjected to CE-MS/MS. Two DIA methods, AIF and vDIA, were investigated in this study. Because a tricky acquisition method for vDIA was employed, we updated the function of the MS-DIAL chromatogram deconvolution function to deal with the acquired data. A total of eight precursor windows, determined by the density of the detected peaks in the *m/z* dimension when analyzing the macrophage cells, were divided into two parts (**Figure S2** and **Table S2**): Part A and Part B. This bifurcation was interspersed with full MS scans in a recurrent sequence. In other words, starting with a Full MS scan, it proceeds to Part A of the vDIA sequence, returns to a full MS scan, transitions to Part B of the vDIA sequence, and then reverts to another full MS scan. This sequence persists in a repetitive manner (see the detailed description in the legend to **Figure S2**). The division of vDIA MS2 sequences was performed to keep the cycle time of the survey scan shorter and the precursor isolation window as small as possible. The migration times of each sample were corrected using the reference MT information from the internal standards. Importantly, the MT correction refers to the shift of data points in each of the CE-MS raw data, which is distinguished from the function of peak alignment, whose function is used to integrate multiple samples’ data. Furthermore, we predicted the migration times for metabolites recorded in the spectral libraries (total 44,257 metabolites) and the MS-FINDER structure database (total 446,485 metabolites) to increase confidence in the annotation process. In fact, the MT information was linearly shifted based on the information from the internal standards detected in the biological samples.

**Figure 1.**
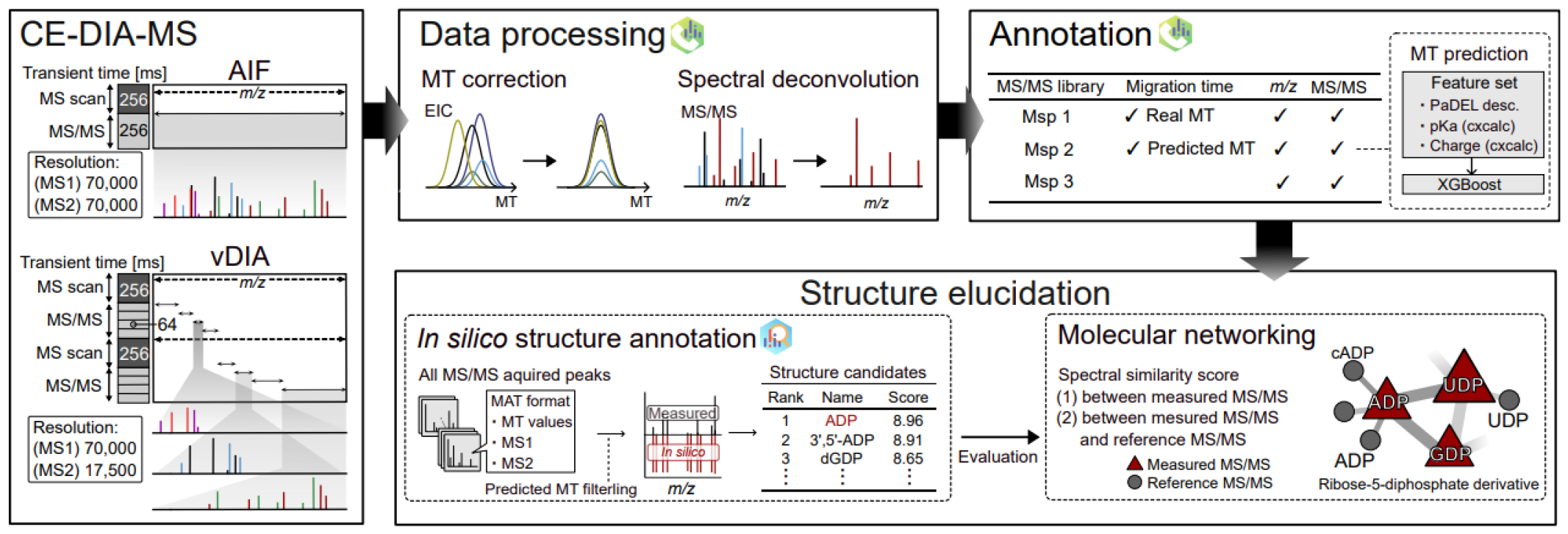
Workflow of CE-DIA-MS/MS for untargeted metabolomics. Comprehensive MS/MS spectra are obtained by two DIA methods, AIF and vDIA. MS-DIAL is used for MT correction using internal standards, MS/MS spectral deconvolution, and annotation at a high confidence level based on predicted MT information and the MS/MS spectra. Furthermore, unannotated peaks are subjected to *in*-*silico* annotation using MS-FINDER, whose results are integrated into the nodes of molecular spectrum networking based on the spectral similarity.

Finally, we prepared three MSP spectral libraries (see **Materials and Methods**): “msp1” contains metabolites having experimental MT values acquired in our laboratory; “msp2” contains metabolites whose MT values were predicted by the machine learning method; and “msp3” contains metabolites with no MT information. We also performed *in-silico* fragment annotation using MS-FINDER with the predicted MT values to annotate the structures of the unannotated compounds. The MS-FINDER outputs were integrated into a molecular spectrum network based on the similarity scores of the MS/MS spectra of metabolite features.

### Preparation of predicted migration time library to improve annotation accuracy

Metabolite annotation is a challenging process in untargeted metabolomics. While the MS/MS spectrum is the most important information, the migration time is also a crucial diagnostic criterion in annotation^2,24,25^. On the other hand, analyzing all the chemicals included in spectral databases in a laboratory is impractical. Thus, we predicted the MT values using an optimized machine learning model. We developed MT prediction models for cation and anion analysis independently because their CE conditions are completely different. The MT values of 267 and 216 metabolites, and 1,306 and 1,400 descriptors were utilized for the cation and anion modes, respectively. The chemical descriptors were generated using PaDEL and ChemAxon. ChemAxon was used to calculate the estimated pKa and charge state of molecules under specific pH conditions, as previous studies have reported that the charge state and molecular size are the most important variables for MT prediction^2,25,26^. We used the Retip R package program to select the best machine learner, and XGBoost was finally selected. The root mean squared error, R-square, and 95% confidence interval for the cation model in the test set were 0.93 min, 0.90, and 1.46 min, respectively, while they were 1.12 min, 0.58, and 1.93 min for the anion model (**Figure 2** and **Table S4**). The top three important variables were charge distribution, bpKa1 (basic pKa), and formal charge for the cation mode, and *m/z*, AATS5e (Average Broto-Moreau autocorrelation - lag 5 / weighted by Sanderson electronegativities), and charge distribution for the anion mode (**Figure S3**). The *m/z* value was calculated by using the exact mass value and the value of the predicted formal charge from ChemAxon. We employed the XGBoost models to predict the MT values for 27,041 and 19,110 compounds from the cation- and anion-MS/MS libraries, respectively. Considering the 95% confidence interval (CI) between experimental and predicted MT values in each model, annotation candidates were filtered with the MT tolerances of 1.46- and 1.93 min using MS-DIAL in cation and anion analysis, respectively.

**Figure 2.**
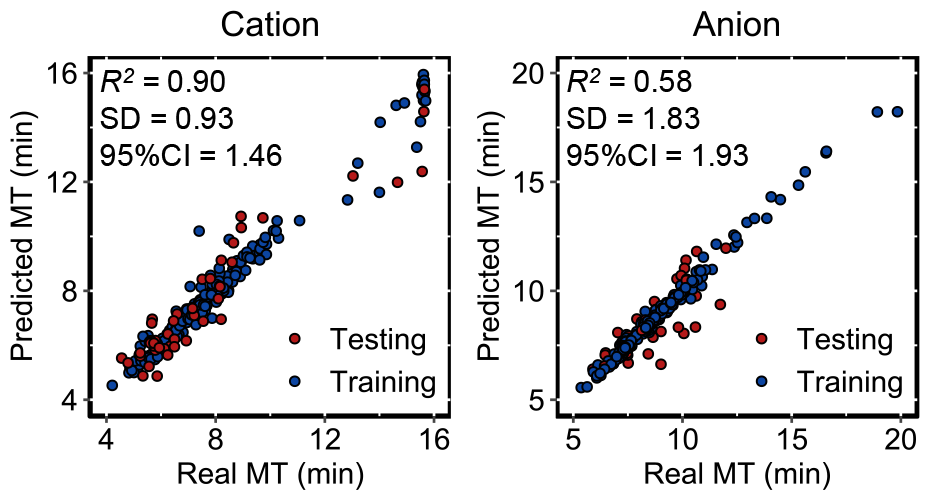
Result of migration time prediction models. The x- and y-axis represent the experimental and predicted migration times (MT), respectively. The MT information of a total of 245 metabolites for cation analysis (left panel) and 180 metabolites for anion analysis (right panel) is used for machine learning. For the test set, 48 and 35 metabolites from the training dataset are used for the model evaluation in cation- and anion analyses. The result of XGBoost is described (see the results of Random Forest in Supplementary Information). Furthermore, the values of R-square, standard deviation (SD), and confidence interval (CI) were described.

### Migration time correction and mass spectral deconvolution for DIA-based identification

The migration times of metabolites are less stable when compared to conventional reverse-phase LC because EOF is affected by temperature, separation buffer volume, and aging of the capillary inner wall^11,12,13,27^. For untargeted metabolomics using CE, these fluctuations hamper the accuracy of peak alignment and metabolite annotation. In this study, we evaluated the retention time correction function of MS-DIAL, where the data points were shifted by the MTs of the internal standards. After detecting the internal standards in all samples, each electrogram data point was corrected by linear interpolation between the two internal standards based on the difference between the measured MT and the reference MT. The MT drifts, as measured by the coefficient of variation (CV) of peaks identified using authentic standard compounds, showed significant improvement with MT correction compared to the original drifts (**Figure 3a**). As a result, the average MT shift was less than 0.2 min, as judged by the results of a total of 76 known compounds, where the maximum peak shift was 0.36 min while the max shift was 1.74 min without MT correction. Peak annotation was also performed using the corrected MT in both cation and anion analyses.

**Figure 3.**
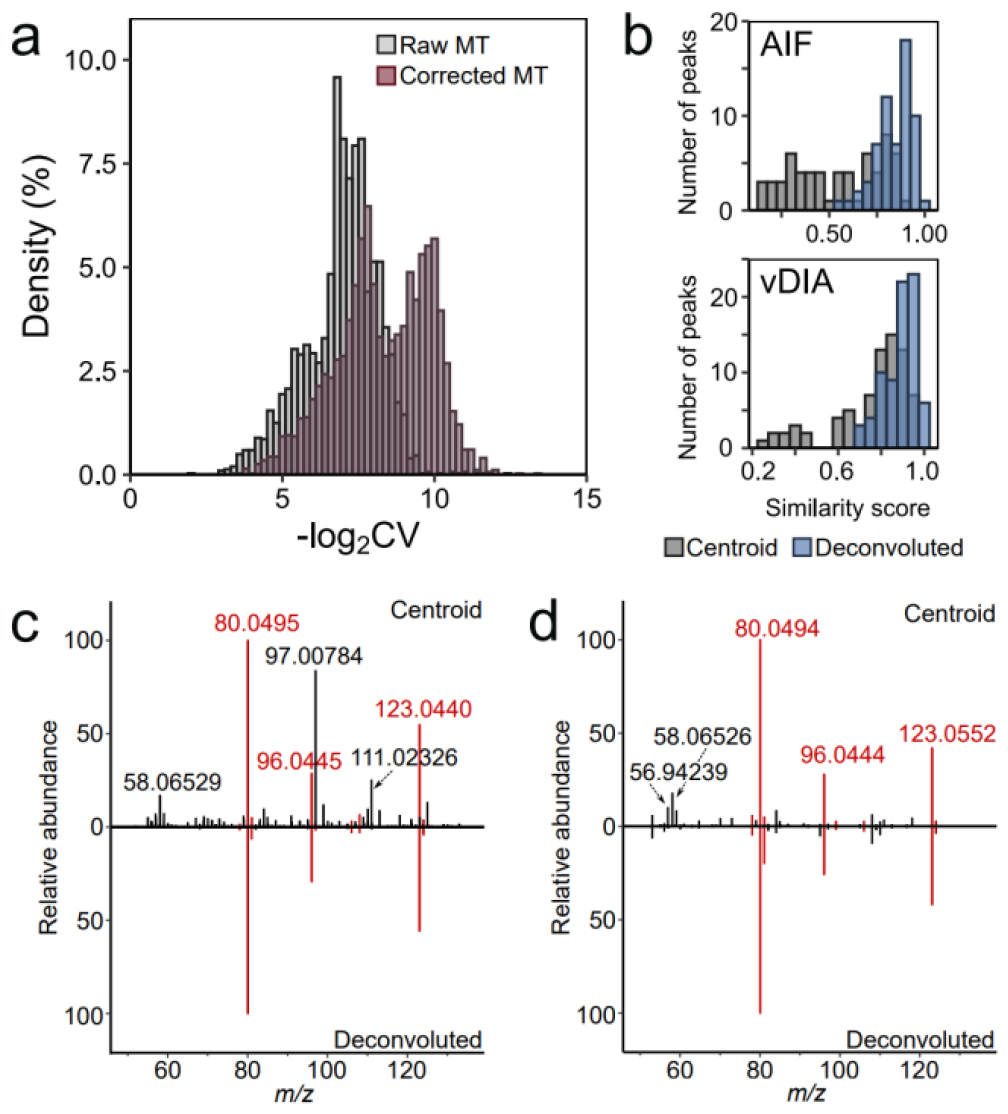
Results of migration time correction and spectral deconvolution for CE-DIA-MS/MS data. (a) Coefficients of variation (CV) before and after the MT correction. The CV values are calculated using the MT table derived from the MS-DIAL program, where the data matrix contains the MT value of each peak of each sample. (b) Similarity score (dot product) distribution of centroid and deconvoluted spectra in AIF and vDIA analysis, where the spectrum was matched with the reference spectra of 63 and 76 metabolites annotated as level 1 confidence in AIF and vDIA mode data. The distribution of similarity score between the experimental spectrum and the reference spectrum is described. (c), (d) Comparison of MS/MS spectra of niacinamide between the centroid and deconvoluted spectra in AIF (c) and vDIA (d) analysis, respectively.

Although MS/MS has rarely been used in CE-MS/MS-based untargeted metabolomics, the MS/MS spectrum is the most important diagnostic criterion for annotation. In fact, annotation results without using MS/MS provided false positives for 3.9% of the characterized peaks estimated in our experimental data with the information of standard compounds, emphasizing that comprehensive MS/MS acquisition is essential for more accurate metabolite annotation (**Table S5**). DIA has great potential for annotation because the MS/MS spectral information can be assigned to all precursor ions, while the MS/MS contamination from different chemicals isolated by the wider precursor window makes the similarity of spectral pattern matching, an important criterion for metabolite annotation, smaller. Thus, we utilized the MS-DIAL chromatogram deconvolution algorithm of MS-DIAL to extract the pure spectrum from such mixed spectra. The results from the two DIA techniques, AIF and vDIA, were evaluated with respect to the spectral matching scores and the annotation rates. The matching scores were substantially improved in both methods when compared to the scores obtained using the non-deconvoluted spectra, i.e., the original centroid spectra (**Figure 3b, 3c and 3d**). Importantly, the improvement, even in the AIF, can be attributed to the high peak capacity of the CE. In fact, the chromatogram deconvolution method in our previous study using AIF in LC-MS/MS did not work well, leading to the development of different deconvolution methods, such as CorrDec^28^. Nevertheless, we observed more improved spectral matching results and annotation rates using CE-AIF-MS/MS. The advantage of AIF is that it decreases the cycle time and increases the accumulation time, contributing to an improvement in peak quantification using MS2 chromatograms. On the other hand, comparing vDIA and AIF, vDIA showed higher spectral matching results and annotation rates owing to its optimized isolation windows (**Table S6 Figure S4e**). Consequently, we decided to use the vDIA data for further analyses, as described below.

### Metabolome analysis of RAW264 cells by CE-vDIA-MS/MS

The metabolic profiling of RAW 267.4 cells after LPS stimulation was performed using CE-vDIA-MS/MS, MT correction and annotation using the predicted MT information. A total of 170 unique metabolites were characterized, of which 76 metabolites were assigned as Metabolomics Standards Initiative level 1 confidence; 40 metabolites were annotated by the precursor *m/z*, predicted MT, and MS/MS matching; and 54 metabolites were annotated by the precursor mass and MS/MS matching (**Table S7**). The metabolome table contains sugar phosphates in glycolysis and the pentose phosphate pathway (PPP), organic acids of the tricarboxylic acid (TCA) cycle, and nucleic acid metabolites, indicating that CE-vDIA-MS enables comprehensive profiling centered on intracellular energy metabolism (**Figure S5** and **Table S7**). Additionally, the separation of isomers (e.g., glucose 6-phosphate, glucose 1-phosphate, isocitrate, and citrate) could be achieved with CE-MS, resulting in high-level annotation based on the precursor *m/z*, MT, and MS/MS. To evaluate the accuracy of the newly annotated compounds based on the predicted MT library, in cation analysis, both the AIF and vDIA showed that three of the newly annotated compounds (3-aminoadipic acid, 2-thio PAF(1-*O*-hexadecyl-2-deoxy-2-thio-*S*-acetyl-sn-glyceryl-3-phosphorylcholine), *N*-acetylneuraminic acid) were confirmed by the authentic standards (**Figure S6**). The result revealed that all compounds were detected within the ±95% Confidence Interval (CI) of the predicted MT, indicating the accuracy of the annotation based on the predicted MT library.

Out of 170 metabolites annotated in vDIA analysis, a total of 111 metabolites were significantly changed after 24 h from LPS administration, including glycolysis metabolites (glucose 6-phosphate, glucose 1-phosphate, and fructose 1,6-bisphosphatase), PPP metabolites (ribulose 1,5-bisphosphate [RuBP], ribose 5-phosphate [R5P], and 2-phosphoglyceric acid [2-PG]), TCA cycle metabolites (citrate, iso-citrate, *cis*-aconitate, and malate), and urea cycle metabolites (citrulline, arginine, creatine, and ornithine) (**Figure 4a and Table S7**). In fact, the terms of these metabolic pathways were enriched in the metabolite set enrichment analysis of MetaboAnalyst^5^ importing significantly altered metabolites information (**Figure 4b**). These metabolic alterations have been reported in LPS-stimulated macrophages owing to the activation of arginine metabolism, aerobic glycolysis, and dysfunction of mitochondrial respiration^29,30,31,32^. Additionally, we observed a significant increase in 2-hydroxyglutaric acid and itaconic acid, known anti-inflammatory immune products that accumulate in inflammatory macrophages^31,32,33^. These results indicate that our untargeted metabolomics procedure works and provides results comparable to those of previous biological studies.

**Figure 4.**
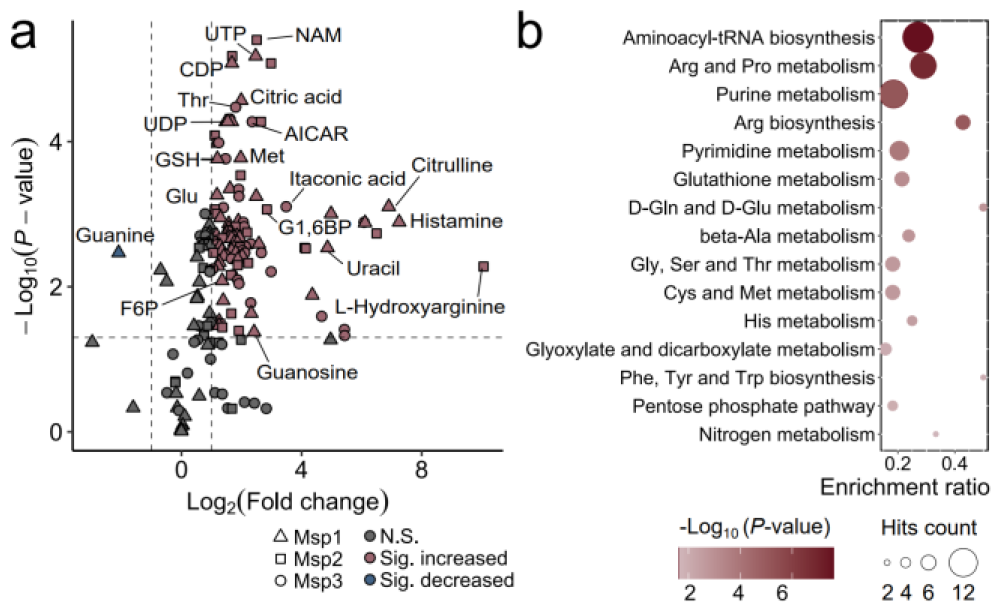
Significantly changed metabolites and enriched metabolites set in between LPS- and control groups. (a) Volcano plot describing significantly changed metabolites in RAW264.7 cells following 24 h of LPS stimulation. Metabolites with more than a 2-fold change and less than a 0.05 adjusted *P* value corrected by false discovery rate in the student t-test were defined as significantly changed metabolites. Node shape denotes the annotation level. (b) Result of Metabolite Set Enrichment Analysis using MetaboAnalyst where significantly changed metabolites were investigated with the KEGG metabolite sets.

Moreover, our data revealed an increase in the levels of spermidine and spermine, referred to as polyamines, in macrophages 24 h after LPS stimulation (**Figure S5**). Spermidine and spermine are cationic polyamines, and several previous studies have reported their pharmacological activities, including antioxidant and anti-inflammatory benefits^31,32,33,34^. Incubation with spermidine before LPS stimulation has been shown to inhibit the production of pro-inflammatory mediators such as nitric oxide (NO) and prostaglandin E2, as well as cytokines including tumor necrosis factor-α and interleukin (IL)-1β in RAW264.7 cells^34^. Additionally, the suppression of IL-6 and IL-1β due to efferocytosis has been demonstrated to be driven by the augmentation of import and accumulation of spermidine and spermine^35^. Our results suggest that the accumulation of polyamines plays a crucial role in orchestrating the transitions from a pro-inflammatory state to an anti-inflammatory state. Consequently, our results suggest that metabolome profiling using CE-vDIA-MS holds the potential to offer novel biological insights and provides more detailed and comprehensive quantitative information on polar metabolites, particularly those associated with energy metabolism.

### Annotation for significantly changed unknowns using MS-FINDER and molecular spectrum networking techniques

We used an *in-silico* structure elucidation tool, MS-FINDER^17^, with the predicted MT values to annotate unknown metabolites. In fact, a total of 1,367 and 1,860 peak features remained unknown in the cation and anion data, respectively. The Human Metabolome Database (HMDB) ^5^ was selected as the structure database, containing 20,805 and 36,871 compounds for cation and anion metabolome data, respectively. The annotation accuracy was evaluated by using the MT- and MS/MS spectral information of known compounds containing 47 peaks for cations and 29 peaks for anions, respectively, resulting in top hits for 64.6% and 65.5% queries, hits within the top three candidates for 79.2% and 79.3% queries, and no candidates for 8.3% (three peaks) and 10.3% (three peaks), respectively (**Figure S7a**). Consequently, we assigned 539 and 611 metabolite names to the unannotated peaks in the cation and anion modes, respectively. Among these assignments, 210 and 258 metabolites were absent from the in-house MT library and MS/MS spectral libraries for cation- and anion analyses, respectively (**Table S8**). The metabolite information assigned by *in*-*silico* structure elucidation should be validated by measuring standard samples. However, on the other hand, complete confirmation using standard compounds for all metabolite candidates is impractical. Therefore, the structural candidates should be evaluated using orthogonal approaches. Here, we constructed molecular networks based on the similarity among the measured MS/MS spectra. In this study, each of the experimental spectra was also compared to the reference product ion spectra included in the spectral libraries **(see Material and Method**). Moreover, the nodes in the molecular networking were further emphasized by the peak intensity and its significance between the control- and LPS-treated groups. As a result, 2,969 peaks were linked by 4,543 edges with a Bonanza score threshold of 0.80 and a maximum edge number per node of 5 (**Figure 5a)**.

**Figure 5.**
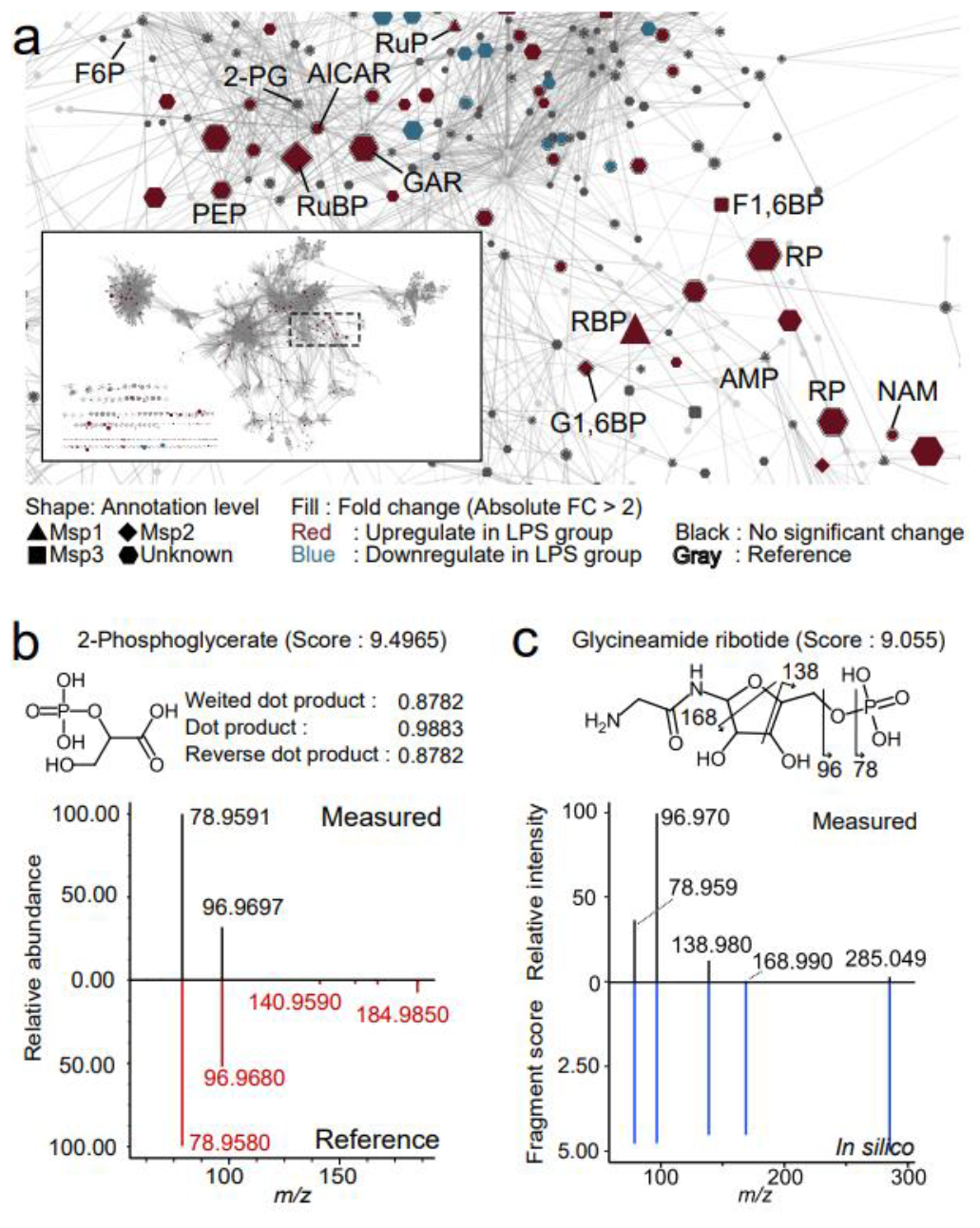
Molecular spectrum networking result from the anion metabolome data. (a) The zoom-in molecular network containing the nodes related to monophosphate ester metabolites. The red, blue, black, and gray colors of the node mean a 2-fold increase in the LPS group, a 2-fold decrease in the LPS group, smaller than a 2-fold change, and a query of reference spectra. The node size represents the absolute log_2_ fold change. Node shape and edge width denote the annotation level and Bonanza score, respectively. The diamond node shape means unknown. The border color becomes black if the peak is annotated by MS-FINDER. The global view of molecular networking is showed in the bottom left. (b) Measured and reference MS/MS spectra of the peak annotated as 2-PG by MS-FINDER. The peak is not characterized, i.e., recognized as a false negative, by searching spectral libraries with the MT filtering due to the number of matched spectra. (c) Annotation of unknown MS/MS spectrum, characterized as glycineamide ribotide (GAR) by checking the literature mining, with the result of the MS-FINDER program.

The algorithm of molecular spectrum networking aims to cluster metabolite ions with similar MS/MS spectral patterns (**Figure 5a**). In particular, metabolites from glycolysis (e.g., glucose 6-phosphate, glucose 1-phosphate, fructose 6-phosphate), the pentose phosphate pathway (PPP) (e.g., 6-phosphogluconic acid, ribulose bisphosphate, xylulose 5-phosphate), and nucleotide metabolism (e.g., cytidine-5’-monophosphate, adenosine monophosphate) were grouped together. This grouping was attributed to their shared structural features, providing fragment ions related to the phosphate groups (**Figure 5a**). For the peaks identified in CE-vDIA-MS, we annotated F6P and ribulose 5-bisphosphate based on the reference MT values, while nicotinic acid mononucleotide (NAM) was annotated using the predicted MT values. Notably, this cluster contained unknown peaks annotated as 2-phosphoglyceric acid (2-PG), phosphoenolpyruvic acid (PEP), and glycineamide ribotide (GAR) by using MS-FINDER (**Figure 5a**). PEP and its substrate 2-PG were directly interconnected, and PEP was upregulated in the LPS group, consistent with the accumulation of metabolites constituting PPP in proinflammatory macrophages (**Figure 4b**). Our measured MTs matched those in our in-house library within a 1 min deviation, and the MS/MS spectra were partially concordant with the reference library’s spectra (**Figure 5b and S7b**). However, their experimental MS/MS spectra often had fewer or less intense product ions, falling below the annotation threshold set when searching spectral libraries. Our results indicated that *in silico* structure annotation tools, such as MS-FINDER, can be used to minimize false negatives and propose structures not recorded in spectral libraries.

Finally, the annotation of glycineamide ribonucleotide (GAR) is showcased as the result of the integrated analysis of MS-FINDER and molecular spectrum networking. The GAR MS/MS data were not recorded in any of the mass spectral libraries. The MS/MS spectrum was linked with those of F6P, G6P, Rl5P, and NAM and increased in the LPS group compared to the control group. The acquired spectrum contained product ions originating from phosphate groups (*m/z* 79 and 96) and phosphate with a segment of ribose (*m/z* 138), and the spectrum is annotated as GAR in MS-FINDER, with a cumulative score of 9.055 out of 10 as the maximum score (**Figure 5e**). Fortunately, we found literature showing the product ion spectrum of GAR derived from collision-induced dissociation using the positive-mode electron-spray ionization method^38^. The fragment pattern in our cation data was similar to that of the literature’s MS/MS spectrum, where diagnostic ions of *m/z* 117, 153, 171, and 251 were detected in both datasets, indicating that the confidence of GAR annotation by MS-FINDER was confirmed by the diagnostic ions of phosphate and ribose in the negative ion mode and by comparing the positive ion mode MS/MS spectrum with that of the authentic standard in the literature (**Figure S7c**). GAR is an intermediate in the purine *de novo* synthesis (PDNS) pathway and further reacted by glycinamide ribonucleotide formyl transferase (GARFT)^39^, which is a potential target for cancer therapy because GARFT activity is linked to cellular proliferation and differentiation. Our metabolome data also showed that other intermediates of the *de novo* synthesis pathway (AICAR, IMP, GMP, GDP, GTP, AMP, ADP, and ATP) were significantly increased in the LPS inflammation group, suggesting that purine metabolism was upregulated via *de novo* synthesis following LPS stimulation (**Figure 4a, 4b, and S5**). Furthermore, the inhibition of purine biosynthesis upon GARFT inhibition has been shown to suppress LPS-induced expression of proinflammatory cytokines in RAW264.7 cells^40^. Our findings suggest a potential association between GAR accumulation and proinflammatory mechanisms in macrophages. Our results indicate that the spectral data from CE-DIA-MS are applicable to *in*-*silico* structure prediction, enabling the exploration of novel metabolic changes without prior MS/MS information.

## CONCLUSION

Our study yields several crucial findings that enhance the utility of CE-MS/MS in untargeted metabolomics. The chromatogram deconvolution algorithm performed effectively for both AIF and vDIA spectra, benefiting from the high peak capacity of CE. Moreover, the annotation rate and spectral matching scores achieved with vDIA surpassed those obtained with AIF. The MT correction provides a framework for aligning multiple CE-MS data, with the MT tolerance parameter set at 0.2∼0.5 min, which is comparable to reverse-phase LC-MS-based large-scale metabolomics data. Additionally, we developed a model for predicting migration time to enhance confidence in metabolite annotation. The proposed workflow for CE-MS/MS untargeted metabolomics was applied to profile the hydrophilic metabolome of macrophage cells in an LPS-induced inflammation model, resulting in the annotation of 170 polar metabolites primarily related to intracellular energy metabolism. Through the integrated approach of MS-FINDER and molecular spectrum networking, we achieved two significant outcomes: 1) the identification of false-negative metabolites initially overlooked in MSP-based annotations and 2) the discovery of a biologically important metabolite, GAR, whose spectrum is absent in any spectral libraries. Utilizing the CE-vDIA-MS approach, we uncovered metabolic alterations in *de novo* purine synthesis, complementing previously reported shifts in glycolysis, the TCA cycle, and arginine metabolism. The results of this study suggest that employing data-independent acquisition (DIA) in CE-based metabolomics allows for global metabolic profiling based on comprehensive MS/MS acquisition and predicted MT values.

## ASSOCIATED CONTENT

### Supporting Information

The Supported Information is available free of charge at xxxxx.

### Data availability

All raw CE-MS data are available on the RIKEN DROPMet website under index number DM0053 (http://prime.psc.riken.jp/menta.cgi/prime/drop_index). The metabolome- and the MS-FINDER annotation results are available as Supplementary Data 1 and Data 2, respectively.

## AUTHOR INFORMATION

Corresponding Author

## Author Contributions

K.O., K.S., H.Y. and H.T. designed the study. Y.M. and H.T. updated the MS-DIAL program. S.K., M.C., T.N., and K.O. prepared the biological samples, and Y.O., T.N., K.S., and H.Y. performed the CE/MS analyses. S.K. performed data analyses. S.K. and H.T. prepared the manuscript. All authors thoroughly discussed this project and helped improve the manuscript.

## ACKNOWLEDGMENT

This study was supported by the JSPS KAKENHI (21K18216, H.T.), National Cancer Center Research and Development Fund (2020-A-9, H.T.), AMED Japan Program for Infectious Diseases Research and Infrastructure (21wm0325036h0001, H.T.), AMED Brain/MINDS (JP15dm0207001, H.T. and H.T.), JST National Bioscience Database Center (NBDC, H.T.), and JST ERATO “Arita Lipidome Atlas Project” (JPMJER2101, H.T.). The study represents a portion of the dissertation submitted by Saki Kiuchi to the Tokyo University of Agriculture and Technology, in partial fulfillment of the requirement for her Ph.D.

---

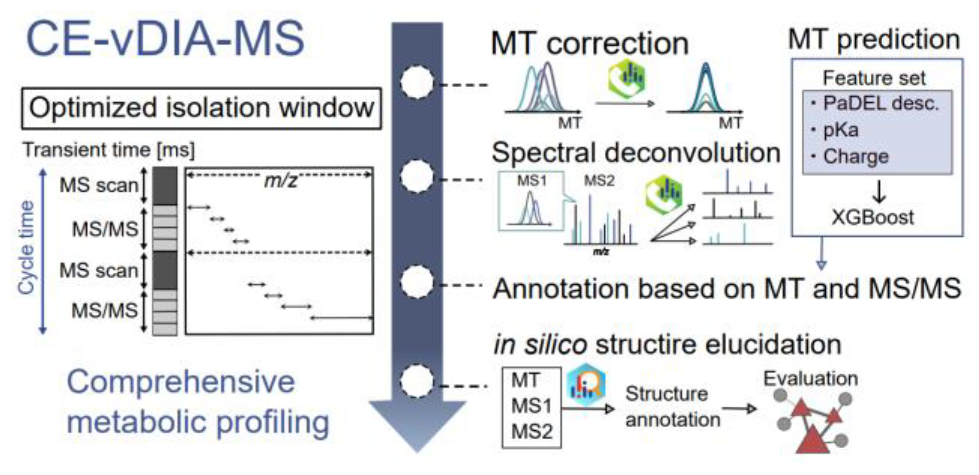

