## Supporting information for "Using variable data independent acquisition for capillary electrophoresis-based untargeted metabolomics"

**Contents**

Figure S1–7

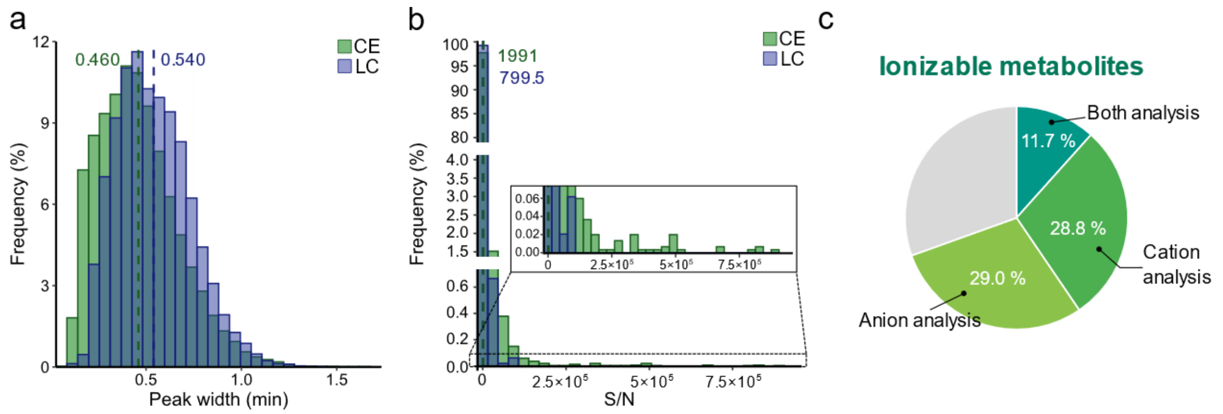


**Supplementary Figure 1. Potential advantages of capillary electrophoresis coupled mass spectrometry (CE/MS)-based untargeted metabolomics.** (a) Statistical analysis of peak widths in conventional CE-MS and reverse-phased liquid chromatography-MS (LC-MS) techniques. The CE-MS and LC-MS data are obtained in this study (refer to Supplementary Information). The x- and y-axes display the peak width (min) and the frequency (%), respectively. The distributions of CE-MS and LC-MS are represented by green and blue colors, respectively, with average peak width values also shown. (b) Statistical analysis of signal-to-noise ratio (S/N) in CE-MS and LC-MS data. The properties in the histograms are explained as in **Figure S1a**. The R package "ggbreak" was utilized to create breaks in the y-axis between 0.6 and 2.0 and between 4.0 and 80. (c) Statistical analysis of cationic- and anionic metabolites under the pH conditions used in CE-MS analysis. A total of 3,727 metabolites, marked as "detected and quantified" in the Human Metabolome Database (HMDB), were considered. The formal charge value of metabolites was calculated using the "cxcalc" function of ChemAxon (http://www.chemaxon.com). Metabolites with positive and negative formal charge values at the defined pH condition were categorized as cationic- and anionic metabolites, respectively.

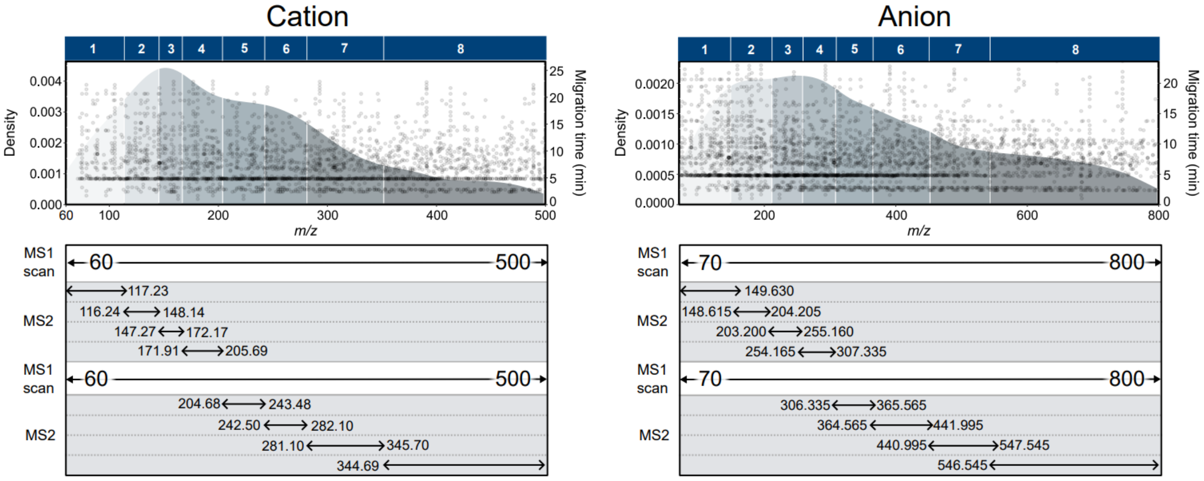


**Supplementary Figure 2. Details of variable data-independent acquisition (vDIA) isolation window settings.** The density plot of detected peaks (top panel) and the optimized vDIA isolation window settings (bottom panel) are depicted for cation (left panel) and anion (right panel) data. Each dot in the density plot represents the migration time (MT) and *m/z* value of detected peaks in CE-MS analysis. The x-axis of the density plot shows the *m/z* value, and the left- and right-y axes represent the frequency (%) and MT value, respectively. The bottom panel illustrates the sequence setting of one scan cycle to parse the entire MS1 and MS2 ranges of interest. For instance, one cycle in cation analysis includes the first full MS1 scanning of *m/z* 60-500, the MS2 scanning of *m/z* 60-117.23 isolation window, and subsequent MS2 scanning of *m/z* 116.24-148.14 isolation window, the MS2 scanning of *m/z* 147.27-172.17 isolation window, the MS2 scanning of *m/z* 171.91-205.69 isolation window, the second full MS1 scanning of *m/z* 60-500, the MS2 scanning of *m/z* 204.68-243.48 isolation window, the MS2 scanning of *m/z* 242.50-282.10 isolation window, the MS2 scanning of *m/z* 281.10-345.70 isolation window, and the MS2 scanning of *m/z* 344.69-500 isolation window. Therefore, the vDIA method used in this study requires two full MS scannings to cover the entire MS2 ranges, defining a "one-cycle" frame in this study.

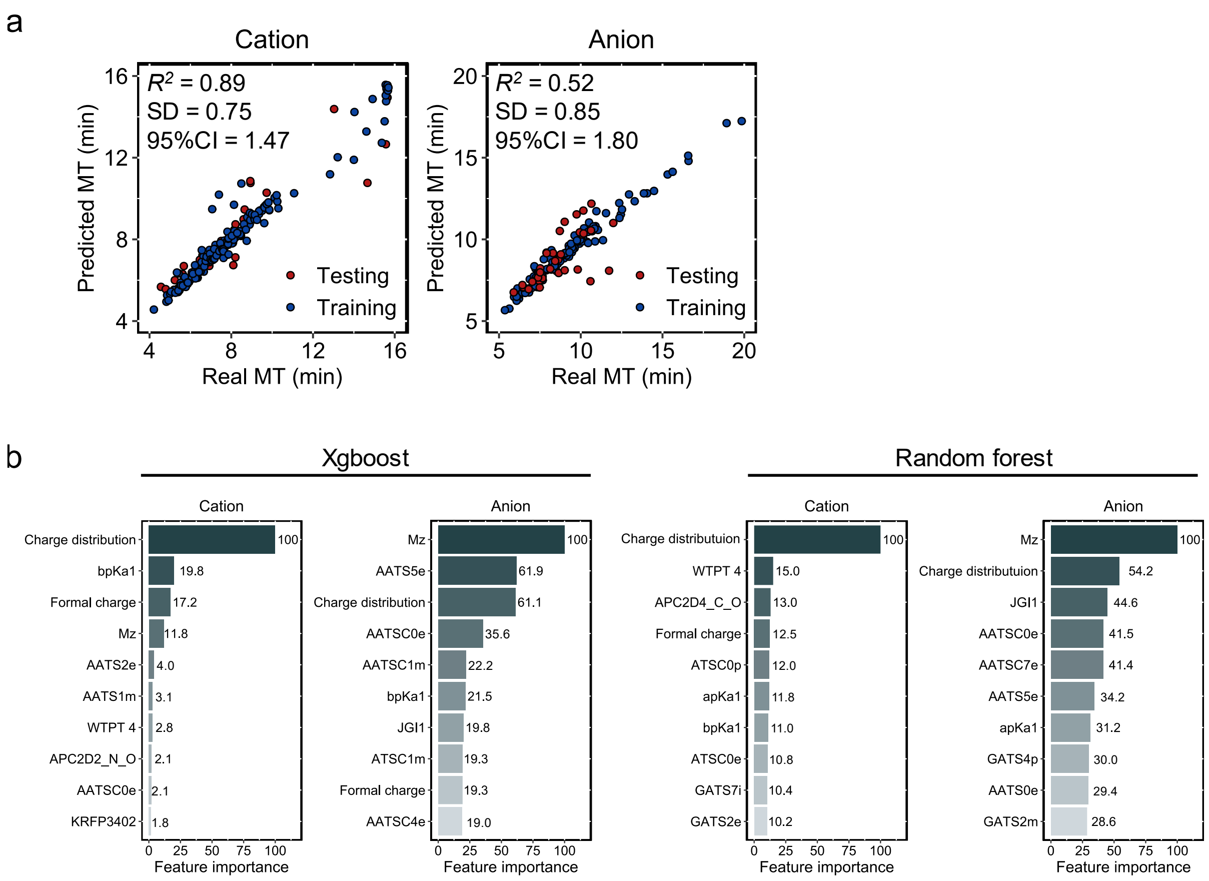


**Supplementary Figure 3**. **Model performances and important variables for migration time predictions (MTs).** (a) Results of MT prediction models using random forest, while the results of XGBoost were described in **Figure 2** of the main text. (b) Top 10 important variables in XGBoost and random forest models, where the largest score was normalized to 100. While the Retip program also evaluates Keras and LightGBM packages, the results of XGBoost and Random Forest outperformed those from Keras and LightGBM programs. Therefore, the details of XGBoost and Random Forest were described.

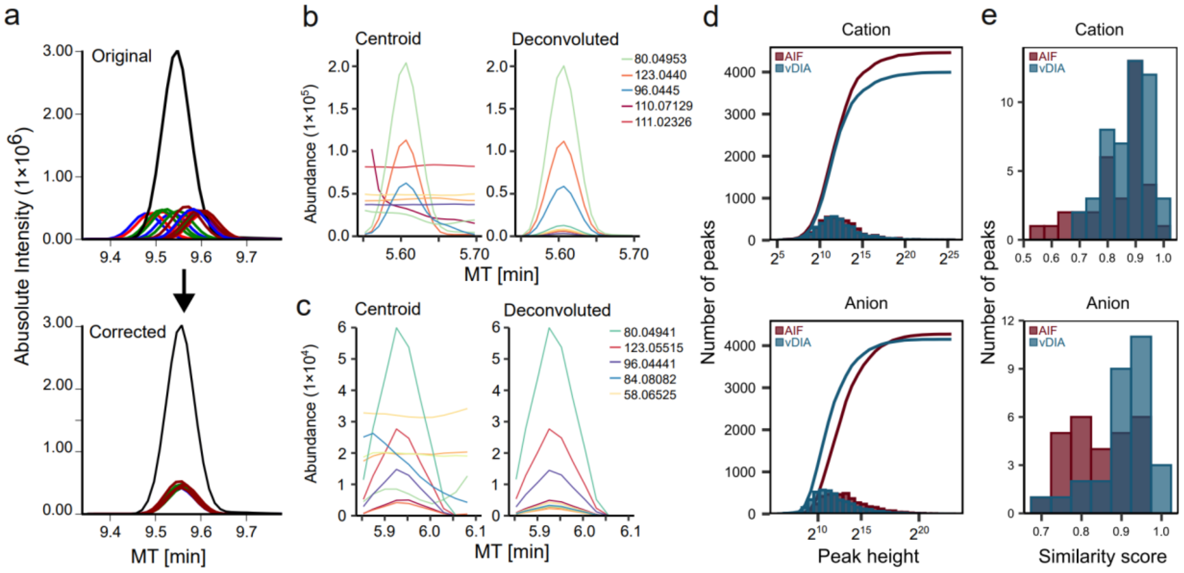


**Supplementary Figure 4. Spectral data comparison between AIF- and vDIA-based untargeted metabolomics.** (a) EIC of an internal standard in cation analysis before and after MT correction. (b), (c) MS/MS chromatograms of niacinamide in AIF (b) and vDIA (c) analysies, respectively. (d) Histogram and cumulative frequency curve of peak heights detected in the pooled QC sample. The x- and y-axis show the peak heights and the number of detected peaks, respectively. Statistics for AIF and vDIA data are described in red and blue, respectively. (e) Comparison of spectral similarity scores between deconvoluted- and authentic standard spectra. Similarity score (dot product) distributions of the cation mode contain the results of similarity matchings of 35 and 48 metabolites for AIF and vDIA analysis, respectively. For the anion mode, scores of 27 and 29 metabolites were described for AIF and vDIA analyses, respectively.

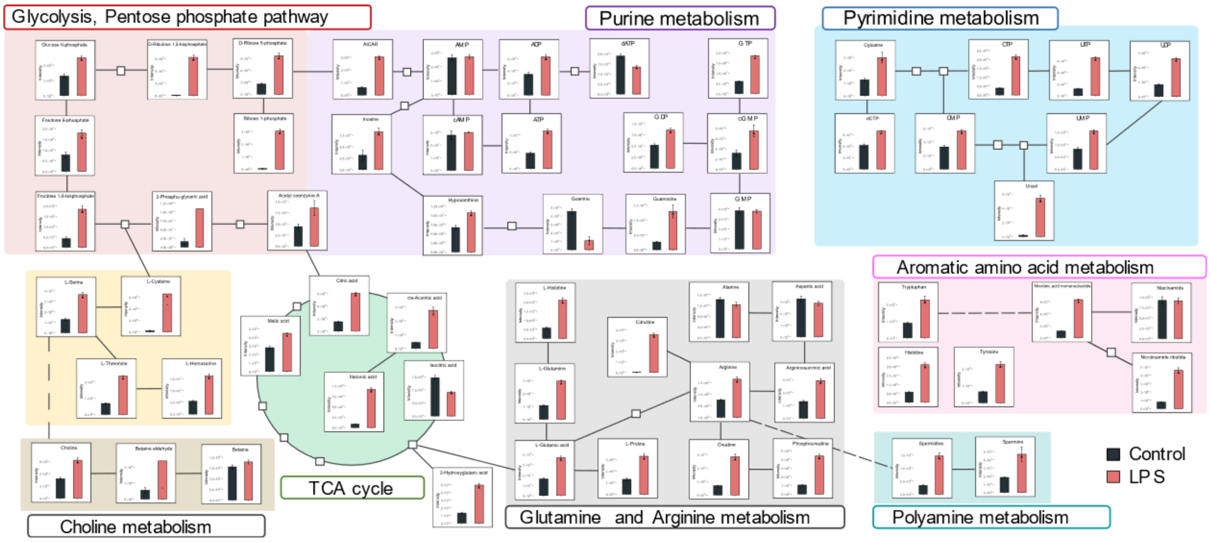


**Supplementary Figure 5. Metabolic profiling results from CE-vDIA-MS-based untargeted metabolomics for LPS-stimulated RAW 264.7 cells.** Summary of the pathway mapping where the major central metabolic pathway is described. Peak height values are normalized by the cell counts and used as the quantitative values. The black- and pink-colored bars represent the control and LPS-group samples, respectively. The bar graph with error bars represents the average +/- the standard error (SE) (n= 4, biological replicates). *$P < 0.05$.

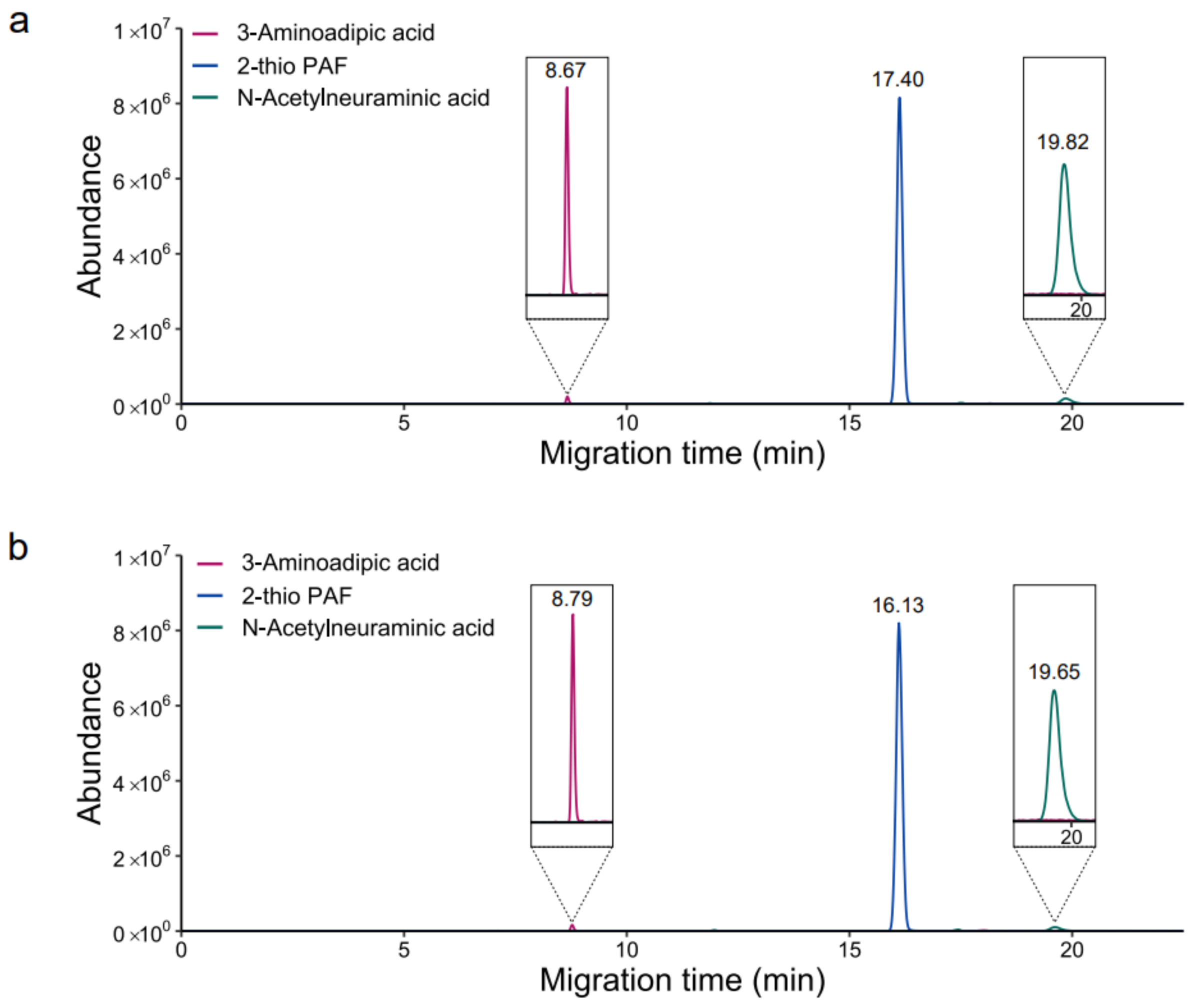


**Supplementary Figure 6. Extracted ion chromatograms (EICs) of 3-aminoadipic acid, 2-thio PAF (1-*O*-hexadecyl-2-deoxy-2-thio-*S*-acetyl-*sn*-glyceryl-3-phosphorylcholine), and *N*-acetylneuraminic acid measured in the cation analysis.** (a) EICs in cellular metabolome. (b) EICs in the authentic standards. The migration times were corrected using the migration times of internal standards.

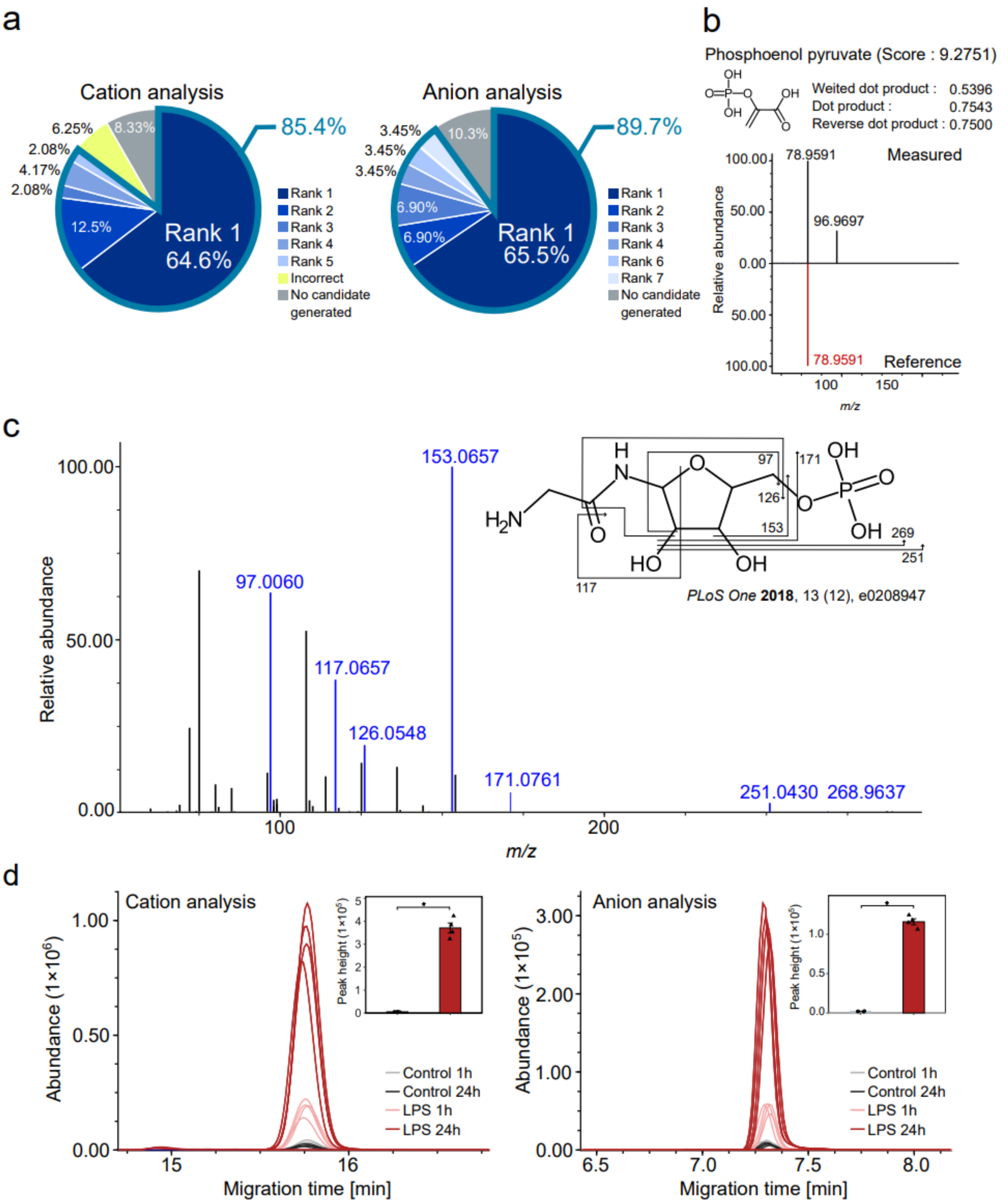


**Supplementary Figure 7. MS-FINDER evaluation in the CE-MS/MS-based untargeted metabolomics platform.** (a) MS-FINDER structure elucidation accuracy with the use of predicted MT data. The spectral queries of 48 and 29 metabolites are imported in cation and anion analyses, respectively, whose annotation could be confirmed by the authentic standard compounds. The candidates are considered as “correct” when the first block of the InChIKey (14 letters) was matched. (b) Measured and reference MS/MS spectra of the peak annotated as phosphoenol pyruvate (PEP) by MS-FINDER. The peak is not characterized, i.e., recognized as a false negative, by searching spectral libraries with the MT filtering. (c) The MS/MS spectrum for a peak annotated as glycineamide ribotide

(GAR). The vDIA spectrum in cation analysis was shown where the blue product ions are consistent with the fragmentation pattern reported by Mádrová et al. (d) Extracted ion chromatograms of GAR in CE-vDIA-MS data analyzing the RAW 264.7 cells. The bar plot shows the normalized peak height of control (gray) and LPS group (red) after 24 hours of LPS stimulation. Error bars represent mean +/- SE (n= 4). *$P < 0.05$.
